# Conserved His-Gly motif of acid-sensing ion channels resides in a reentrant ‘loop’ implicated in gating and ion selectivity

**DOI:** 10.1101/2020.03.02.974154

**Authors:** Nate Yoder, Eric Gouaux

## Abstract

Acid-sensing ion channels (ASICs) are proton-gated members of the epithelial sodium channel/degenerin (ENaC/DEG) superfamily of ion channels and are expressed throughout central and peripheral nervous systems. The homotrimeric splice variant ASIC1a has been implicated in nociception, fear memory, mood disorders and ischemia. Here we extract full-length chicken ASIC1a (cASIC1a) from cell membranes using styrene maleic acid (SMA) copolymer, yielding structures of ASIC1a channels in both high pH resting and low pH desensitized conformations by single-particle cryo-electron microscopy (cryo-EM). The structures of resting and desensitized channels reveal a reentrant loop at the amino terminus of ASIC1a that includes the highly conserved ‘His-Gly’ (HG) motif. The reentrant loop lines the lower ion permeation pathway and buttresses the ‘Gly-Ala-Ser’ (GAS) constriction, thus providing a structural explanation for the role of the His-Gly dipeptide in the structure and function of ASICs.

---

In humans, four ASIC genes, in concert with splice variants, encode for at least eight distinct subunits that assemble as proton-gated, voltage-insensitive heteromeric or homomeric channels. The homotrimeric splice variant, ASIC1a, is found on the dendrites, post-synaptic spines, and cell bodies of central neurons (1, 2) and is enriched in the amygdala (3). ASIC1a channels participate in multiple central nervous system (CNS) processes including fear conditioning (4–6), nociception (7, 8) and synaptic plasticity (9–12). ASICs are also therapeutic targets (13–15), with localization patterns, Ca^2+^ permeability and proton-dependent activation implicating these channels in acidosis-induced neuronal injury (16–19) and mood disorders (20, 21). Upon activation by excursions to low pH, homotrimeric ASIC1a channels open, exhibiting modest Na^+^ selectivity with P_Na_ / P_K_ ~ 7.8 and P_Na_ / P_Ca_ ~ 18.5 (22, 23) and subsequently entering a long-lived desensitized state in hundreds of milliseconds (24). A simple gating mechanism consistent with the observed kinetic measurements is comprised of high pH resting, low pH open and low pH desensitized states (25).

ASICs, and by extension, members of the ENaC/DEG superfamily of ion channels, are trimers, with each subunit composed of large extracellular domains, two transmembrane helices, and intracellular amino and carboxyl termini (26). Chicken ASIC1a is the structurally most well-characterized member of the ENaC/DEG family of ion channels, with x-ray and single-particle cryo-EM structures of the detergent-solubilized channel determined in the resting (27), open (28) and desensitized (26, 29) states. These structures, defined by conducting and non-conducting pore profiles, ‘expanded’ and ‘contracted’ conformations of the thumb domain, and distinct conformations of critical β-strand linkers (26–31), as examples, have provided the foundation for structure-based mechanisms for proton-dependent gating and desensitization. However, in all of these structures the intracellular amino terminal residues Met 1 through Arg 39, including the highly conserved HG motif, are disordered and not visible in the x-ray diffraction or cryo-EM density maps, leaving a significant gap in our understanding of how these regions contribute to channel structure and function.

Numerous lines of evidence indicate that the amino termini of ENaC/DEG channels contribute to gating and ion conduction properties, with this region including the conserved HG motif. Indeed, mutation of the conserved glycine in the β subunit of ENaC channels reduces channel open probability (32) and underlies one form of pseudohypoaldosteronism type 1 (PHA-1) (33). In ASICs, pre-transmembrane domain 1 (pre-TM1) residues participate in ion selectivity and proton-dependent gating (34), further demonstrating that the amino terminus of ENaC/DEG channels plays an important role in ion channel function and may comprise portions of the ion pore. Despite the wealth of structural information surrounding ASICs and, more recently, the ENaC structure (35), the architecture of the cytoplasmic terminal domains as well as the molecular mechanisms by which the HG motif and pre-TM1 residues contribute to channel structure and function have remained elusive.

Here we present structures of cASIC1a solubilized without the use of detergent, using SMA copolymers (36, 37), in distinct conformational states at low and high pH. Our results reveal that amino terminal pre-TM1 residues form a reentrant loop, and that the HG motif is situated ‘below’ the GAS belt TM2 domain swap, at a subunit interface and along the lower ion permeation pathway. Furthermore, we show that the lower half of the ion permeation pathway in resting and desensitized states is comprised entirely of pre-TM1 reentrant loop residues, informing mechanisms for the contribution of the amino terminus to ion permeation properties. Finally, we observe putative lipid density at the transmembrane domain (TMD) that may indicate the preservation of endogenous protein-lipid interactions maintained by detergent-free channel isolation methods. These studies provide structures of an SMA-solubilized ASIC and uncover the architecture of a motif central to gating and ion permeation.

## RESULTS

### Isolation and structure determination of cASIC1a in SMA copolymer

To elucidate structures of cASIC1a bound with endogenous lipids, we extracted and purified recombinant channels in the presence of SMA copolymer. After a two-step chromatographic purification procedure, SMA-cASIC1a particles were ~95% pure as judged by SDS-PAGE and were monodisperse as measured by fluorescence-detection size exclusion chromatography (FSEC) (38) (Supplementary Data Figure 1A-B). Negative stain transmission electron microscopy also demonstrated good particle distribution and limited aggregation (Supplementary Data Figure 1C).

We next pursued single-particle cryo-EM of SMA-cASIC1a, obtaining reconstructions of ASIC1a channels in low pH desensitized (pH 7.0) and high pH resting (pH 8.0) conformations at estimated resolutions of ~ 2.8 and 3.7 Å, respectively, by gold-standard FSC (39) (Figure 1A-B, Supplementary Data Table 1, Supplementary Data Figures 2-5). While analysis of proton-dependent activation and steady-state desensitization curves for cASIC1a support the presence of a narrow window current at ~ pH 7.0 (27), extensive 3D classification of SMA-cASIC1a particles prepared at pH 7.0 did not indicate the presence of either open or resting channels. We therefore speculate that either the solution or the grid preparation conditions result in stabilization of the desensitized state at pH 7.0.

**Figure 1.**
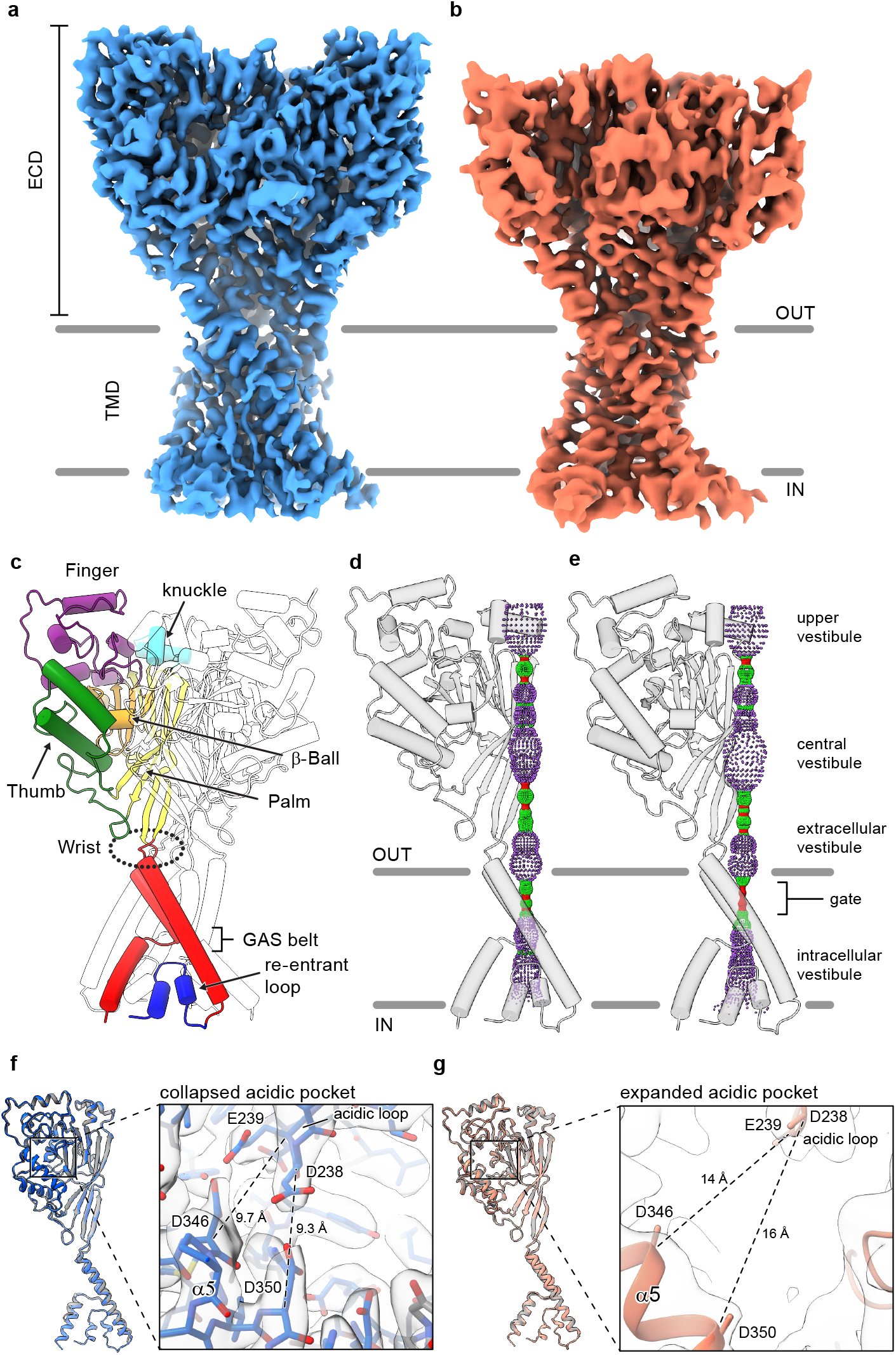
Structures of SMA-cASIC1a. **a-b**, Cryo-EM maps of SMA-cASIC1a at pH 7.0 (**a**) and pH 8.0 (**b**). **c**, Cartoon diagram of cASIC1a with single subunit shown colored by domain. **d-e**, Pore profiles for SMA-cASIC1a in a desensitized (**d**) and resting (**e**) state calculated with HOLE software (pore radius: red < 1. 15 Å < green < 2.3 Å < purple). **g**, Single subunit superposition of SMA-cASIC1a in a desensitized state (blue) and the desensitized state x-ray structure (28, 29) (PDB 4NYK, grey). Detailed view of the collapsed acidic pocket is shown in the inset. **h**, Single subunit superposition of SMA-cASIC1a in a resting state (salmon) and the desensitized state x-ray structure (27) (PDB 5WKU, grey). Detailed view of the expanded acidic pocket is shown in the inset.

In accord with previously solved structures, the homotrimeric ASIC1a channel resembles a clenched fist (26), harboring domain-swapped TM2 helices (28) (Figure 1C). Both the desensitized and resting channels adopt closed ion channel gates, as evidenced by a constriction between residues 433-436 within the upper third of the TMD (Figure 1D-E). At pH 7.0, SMA-cASIC1a particles prepared at pH 7.0 populate a desensitized state that mirrors the overall architecture of the existing x-ray structure (29), including the presence of a proton-bound ‘collapsed’ acidic pocket (Figure 1F). In contrast, SMA-cASIC1a particles maintained at pH 8.0 occupy a high pH resting conformation, characterized by an expanded acidic pocket (Figure 1G) that resembles the high pH, resting state structures solved by x-ray crystallography and single particle cryo-EM (27). We propose that the limited resolution of the resting channel structure, while presumably impacted by sample conditions including thicker ice, may also be due to structural flexibility inherent to the resting channel conformation in the absence of divalent cations, which serve to stabilize an expanded acidic pocket at high pH (40) but which are incompatible with current SMA-based purification strategies.

### Amino terminal residues form a reentrant loop

Numerous experiments have implicated residues within the pre-TM1 region of ASICs and ENaCs in both gating and selectivity (32, 34, 41). Indeed, the highly conserved HG motif is located within the pre-TM1 region of ASICs and ENaCs and its disruption lowers the open probability in ENaCs and underlies PHA type 1 disorder (32, 33). In contrast to existing structures of ASICs solubilized in detergent micelles, we observed strong protein density corresponding to amino terminal residues in cryo-EM maps of both desensitized and resting SMA-cASIC1a channels maintained in a lipid environment (Figure 2A-B).

**Figure 2.**
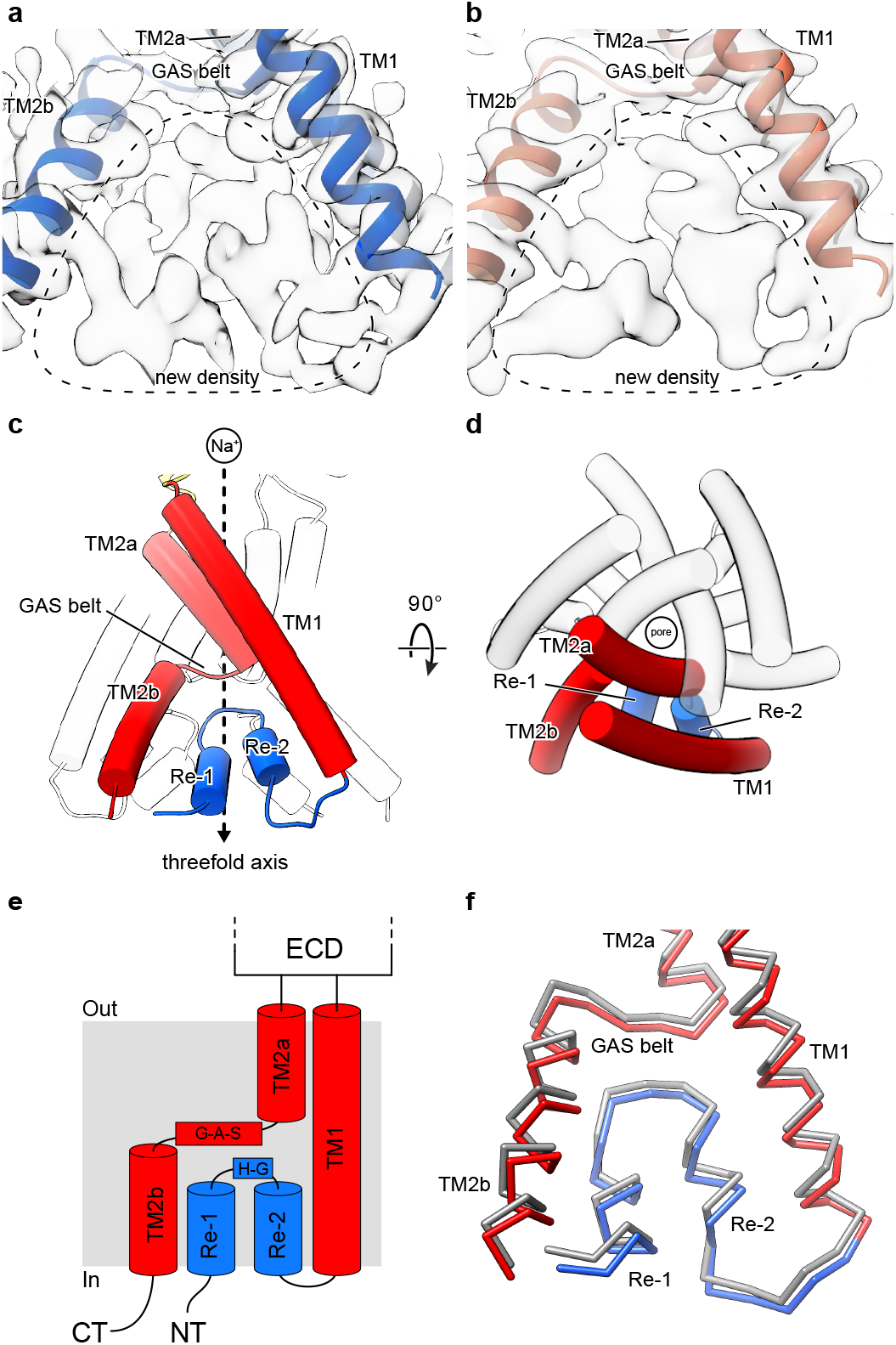
Cryo-EM density and structure of the reentrant, pre-TM1 domain. **a-b**, Cryo-EM density corresponding to amino terminal residues of SMA-cASIC1a at pH 7.0 (**a**) and pH 8.0 (**b**). **c-d**, Side (**c**) and top-down (**d**) views of the TMD from SMA-cASIC1a in a desensitized state at low pH. A single subunit is shown, colored by domain. **e**, Schematic depicting the TMD topology of cASIC1a channels. **f**, Backbone superposition of SMA-cASIC1a in the desensitized (colored by domain) and resting (grey) states.

The quality of the 2.8 Å density map of the desensitized channel was sufficient to model the amino terminus of the existing model (28, 29) (PDB 4NYK) from residue 17 (Supplementary Data Figure 2), demonstrating that pre-TM1 residues form a reentrant loop comprised of two short helical segments separated by a turn, positioned on the cytoplasmic side of the GAS belt (Figure 2C-E). Interestingly, the presence of the reentrant loop does not noticeably impact the position of either transmembrane helix from those observed in prior x-ray or cryo-EM structures. Rather, the reentrant loop residues are ‘pinned’ within the inverted ‘v-shaped’ cavity formed between the lower transmembrane helices and maintained primarily by virtue of intra-subunit contacts with TM2b and TM1 (Supplementary Data Figure 6).

While the quality of the resting channel density map that includes the pre-TM1 residues was not sufficient for unambiguous model building (Supplementary Data Figure 4), no significant differences in reentrant loop conformation were observed between the desensitized or resting channels at the current resolutions (Figure 2F), allowing us to rigid body fit the pre-TM1 structural element derived from the desensitized state structure into the resting state map. In contrast with the discovery of new density for the pre-TM1 region, we did not observe any additional interpretable density associated with carboxy terminus in either the desensitized or resting state maps, thus suggesting that even in SMA-solubilized protein, these regions are disordered.

### The reentrant loop harbors the HG motif

Separated by more than 400 residues in the amino acid sequence, the GAS and HG motifs are highly conserved amongst ENaC/DEG and ASIC channels (Figure 3A) and have been implicated in gating (41) and ion selectivity (42–45). Interestingly, the HG motif, which contains a well-characterized disease mutation at the universally conserved glycine residue (32, 33), is situated on the turn between the reentrant helices where it buttresses the TM2 domain swap and GAS belt residues from ‘below’ (Figure 3B). Residing along the ion permeation pathway and at a subunit interface, the HG motif is capped by the carboxyl terminus of TM2a via an intra-subunit hydrogen bonding interaction with Ile 442 and participates in an inter-subunit hydrogen bonding interaction with a neighboring GAS belt residue via Ser 445 (Figure 3C). This intricate network of intra- and inter-subunit interactions, formed between highly conserved motifs via the TM2 domain swap and amino terminal reentrant loop, underscores the critical nature of the lower pore architecture to ASIC, and by extension, to ENaC function.

**Figure 3.**
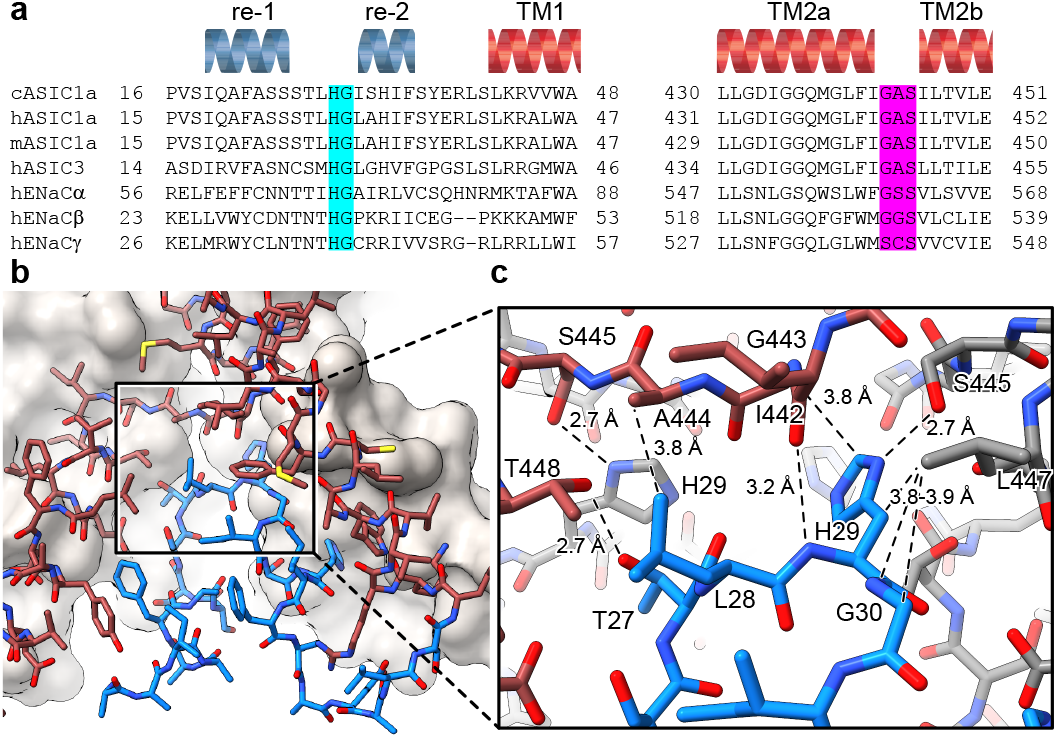
The HG motif resides at a subunit interface ‘below’ the GAS belt. **a**, Sequence alignment of selected ASIC1, ASIC3 and ENaC channels covering the pre-TM1 and TM2 domains with GAS domain and HG motif residues highlighted in pink and light blue, respectively, and with secondary structure for cASIC1a shown above the sequences. **b-c**, View of the chemical environment around the reentrant loop (**b**) with a detailed view of the GAS domain and HG motif interface(**c**).

### Pre-TM1 residues form the lower ion permeation pathway

In structures of both desensitized and resting SMA-cASIC1a channels, the ‘upper’ ion permeation pathway is comprised of TM2a residues and contains a closed gate between Gly 432 and Gly 436, in agreement with existing x-ray and cryo-EM models (Figure 4A). However, where structures of ASIC1a in resting (27), open (28) and desensitized (29) conformations highlight a lower ion permeation pathway comprised entirely of TM2b residues that expands outwards to form a wide intracellular vestibule, pre-TM1 residues of the SMA-isolated cASIC1a channels line a more narrow ion permeation pathway extending below the GAS belt (Figure 4A-B).

**Figure 4.**
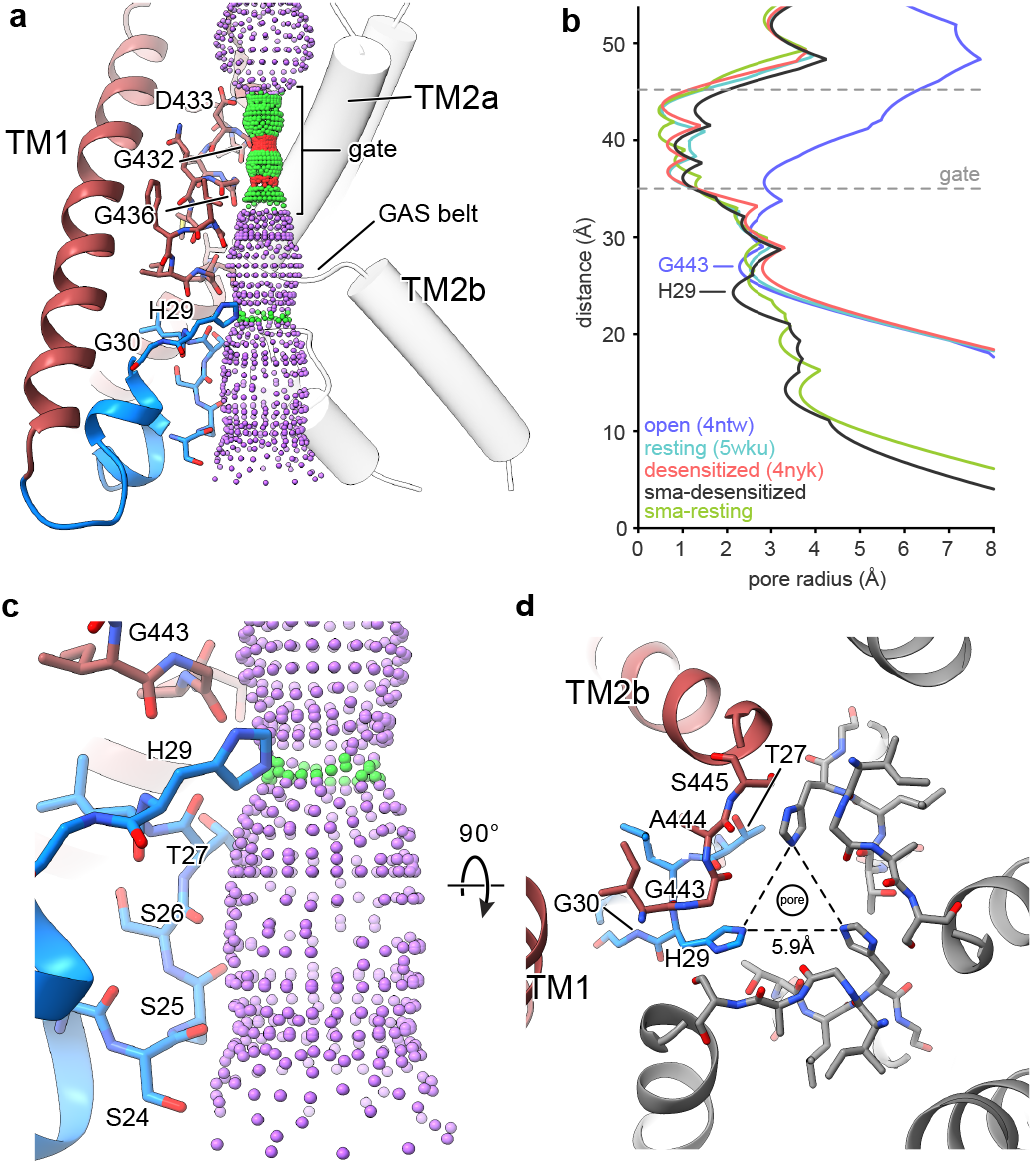
The reentrant loop forms the lower ion permeation pathway. **a**, Overview of pore-forming residues of desensitized SMA-cASIC1a channels beginning at the ion channel gate. Pore profile calculated with HOLE software (pore radius: red < 1. 15 Å < green < 2.3 Å < purple) is shown. **b**, Plot of pore radius corresponding to the view in (**a**) for resting (27) (PDB 5WKU), open (28) (PDB 4NTW) and desensitized (28, 29) (PDB 4NYK) x-ray and SMA-cASIC1a cryo-EM structures. **c**, Detailed view of the ‘lower’ ion permeation pathway formed by pre-TM1 residues. **d**, Top-down view of the constriction ‘below’ the ion channel gate formed by His 29, as visualized in the desensitized SMA-cASIC1a state. The GAS belt residues are in the foreground.

The lower ion conduction pathway of resting and desensitized SMA-cASIC1a channels is formed by reentrant amino terminal residues Ser 24 through His 29 (Figure 4C), the latter of which is situated immediately below the GAS belt and is oriented towards the threefold axis where it forms a constriction below the gate in the desensitized channel (Figure 4D). Our data demonstrate that pre-TM1 residues line the lower ion conduction pathway in structures of resting and desensitized cASIC1a channels, providing a structural rationale for earlier reports which indicated that pre-TM1 residues may form part of the pore (46) and contribute to ion permeation and Na^+^ selectivity of ASICs (34).

In the x-ray structure of an open channel conformation, hydrated Na^+^ ions encounter a constriction at the GAS belt TM2 domain swap (28), which has long been thought to underpin ion selectivity in ENaC/DEG channels (42, 44, 47). Recently, however, residues along TM2a and TM2b both ‘above’ and ‘below’ the GAS belt have been demonstrated to be important determinants of selectivity in ASIC1a (48). Despite the presence of an ordered reentrant loop and a narrower pore, we did not observe a change in the position of either TM2a/b or the GAS belt residues between the resting and desensitized conformations. Additional studies of the open state, perhaps under SMA isolation conditions, will be required to illuminate the structure of the activated, ion conducting state of the channel.

### Density features suggest TMD-membrane interactions

The reconstitution of membrane proteins into lipid nanodiscs is a well-established technique in structural biochemistry that permits the study of sensitive membrane proteins embedded in phospholipid bilayers (49, 50). While a reconstitution approach provides for a controlled and defined lipid environment, the necessity of an initial detergent-based extraction step may disrupt protein-lipid interactions integral to the structural integrity of transmembrane segments. In contrast with nanodisc reconstitution, SMA copolymers extract membrane proteins directly from the lipid bilayer, eschewing detergent entirely and permitting the study of membrane proteins in the presence of endogenous lipids (51, 52) and, in principle, maintaining the protein-lipid interactions that occur at the cell membrane (53).

In our 2.8 Å reconstruction of a desensitized ASIC1a, we observed multiple ordered elongated densities situated in lipophilic channels along the TMD (Figure 5A) that we suggest may correspond to bound lipids. Separated into spatially distinct clusters (Figure 5B), putative lipid densities reside near the top of the membrane sandwiched between TM2a and TM1 helices (Figure 5C) and near the bottom of the membrane between TM1 and TM2b (Figure 5D).

**Figure 5.**
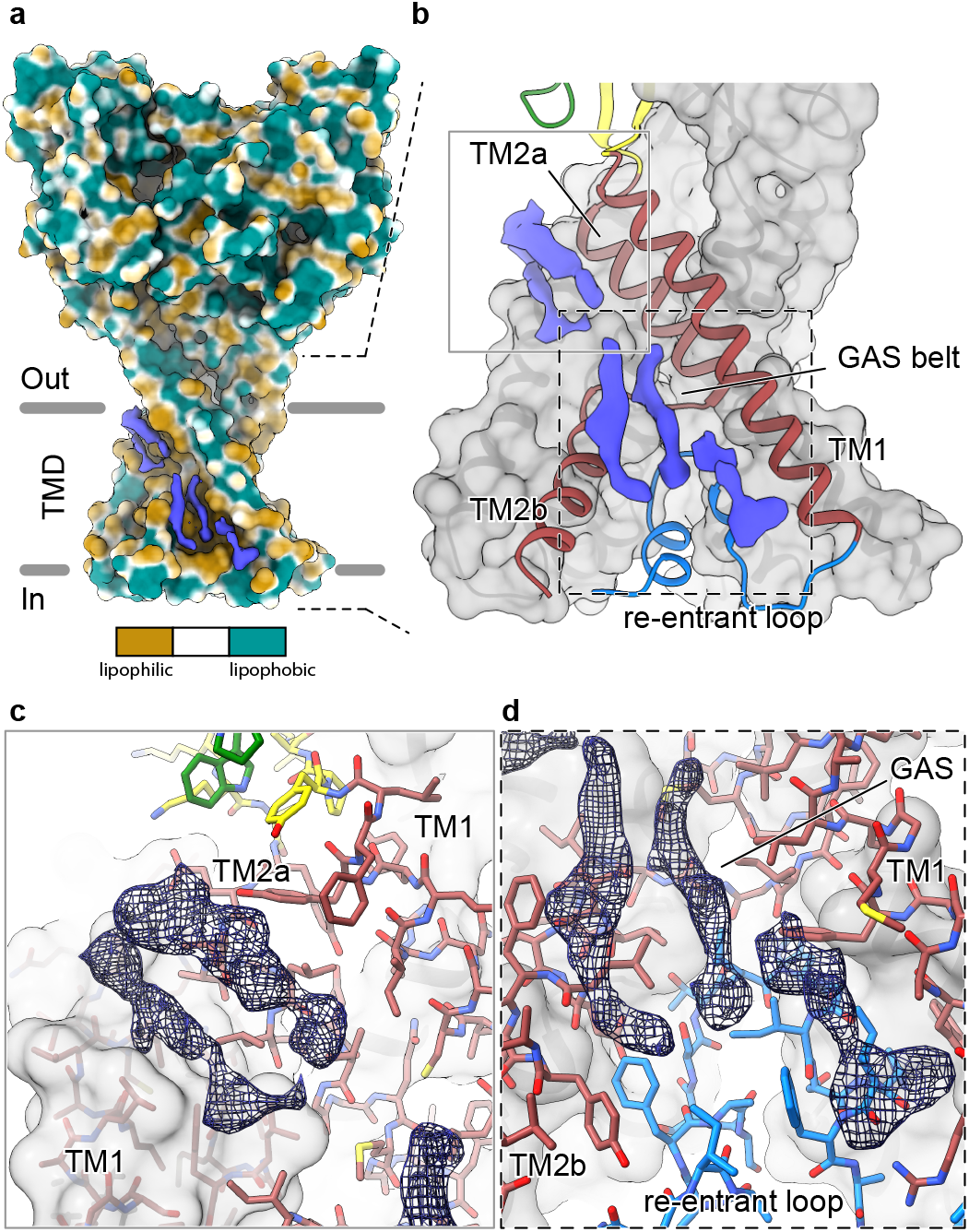
Elongated density within lipophilic channels at the TMD of SMA-cASIC1a channels. **a**, Surface representation of SMA-cASIC1a in the desensitized state colored by lipophilicity potential calculated with pyMLP (60) in ChimeraX (61). **b**, Hybrid cartoon and surface representation of putative lipid sites at the TMD. **c-d**, Putative lipid densities between TM1 and TM2a (**c**) and between TM1 and TM2b adjacent to the reentrant loop (**d**) of SMA-cASIC1a.

Our results suggest that the local lipid environment is important for maintaining the architecture of the reentrant pre-TM1 residues in ASIC1a and thus the integrity of the lower pore pathway. Therefore, the location of at least one cluster of putative lipid densities within a lipophilic cleft immediately adjacent to reentrant loop residues and the GAS belt (Figure 5D) is intriguing, especially given that the cryo-EM structure of a full-length cASIC1a channel in an n-dodecyl-β-D-maltoside lacked ordered amino terminal residues (27). However, given the resolution of our SMA-cASIC1a reconstructions, we are unable to assign this density to any specific lipid. Future experiments are needed to determine the molecular composition of SMA-cASIC1a particles and to explore relevant interactions between ASICs and the plasma membrane.

## DISCUSSION

Here we present structures of chicken ASIC1a solubilized by SMA in high pH ‘resting’ and low pH ‘desensitized’ conformations. While the conformation of both resting and desensitized channels throughout the ECD faithfully mirrors those solved previously via detergent-based methods, our structures demonstrate that amino terminal residues prior to TM1 form a reentrant loop that comprises the lower portion of the ion permeation pathway. In both resting and desensitized structures, the conserved HG motif is situated within the reentrant loop, immediately below the GAS belt TM2 domain swap, where it forms a constriction along the ion permeation pathway and is stabilized by a complex network of inter and intra subunit interactions. Finally, we detected elongated ordered densities within lipophilic channels of the TMD, some of which are adjacent to the reentrant amino terminus, that may correspond to bound lipids.

These results provide a structural basis for contributions of the amino terminus to both ion permeation and proton-dependent gating of ASICs, reveal the location of the conserved HG motif along the ion conduction pathway and expose a role for the plasma membrane in maintaining the TMD architecture of ASICs. Given the structural similarities between ENaCs and ASICs, as well as the highly conserved and functionally-important nature of the HG and GAS belt residues, these results provide detailed structural information pertaining to a pair of motifs central to gating and ion permeation and of possible therapeutic relevance to the entire superfamily of ENaC/DEG ion channels.

## METHODS

### Expression and purification of cASICla channels

Recombinant full-length acid-sensing ion channels (Gallus gallus) were expressed in HEK293S GnTI-cells and membrane fractions were isolated as previously described (27). Membrane pellets were resuspended in ice cold Tris-buffered saline (TBS, 20 mM Tris pH 8.0 and 150 mM NaCl) containing protease inhibitors, dispersed using a dounce homogenizer, and solubilized for 1 hour at 4°C by addition of SL30010 (Polyscope) SMA copolymers to 2% (w/v) final concentration. Membrane debris was removed via centrifugation at 125,171 rcf for 30 minutes at 4°C and the supernatant was incubated with Ni-NTA beads overnight at 4°C in the presence of 10 mM imidazole.

The Ni-NTA bead suspension was then transferred to a XK-16 column and subject to two washes, first with three column volumes of TBS containing 10 mM imidazole and last with three column volumes of TBS containing 30 mM imidazole. The SMA-cASIC1a protein was eluted with TBS containing 250 mM imidazole, and peak fractions were pooled and concentrated to ~ 5 mg/ml. The His_8_ EGFP tag was removed via thrombin digestion using a ratio of cASIC1a to thrombin of 25:1, overnight at room temperature (RT). The following day, the SMA-cASIC1a protein was purified via size-exclusion chromatography (Superose 6 10/300) using a mobile buffer composed of TBS supplemented with 1 mM DTT. A single peak fraction was collected and concentrated to ~ 1 mg/ml for cryo-EM sample preparation.

### Cryo-EM of SMA-cASIC1a

Quantifoil holey carbon grids (R1.2/1.3 200 mesh Au) were glow discharged for 1 minute at 15 mA, carbon side facing up. For structure determination of SMA-cASIC1a particles at high pH, purified protein at ~ 1 mg/ml was used immediately for grid preparation. For structure determination at low pH, the pH of the sample was adjusted to 7.0 by addition of MES, pH 6.0, following concentration of purified protein to ~ 1.0 mg/ml. A 4 μl droplet of sample, applied to the carbon side of the grid, was blotted manually with pre-cooled filter paper (Whatman, grade 1) and the grids were vitrified in ethane/propane mix using a custom-built manual-plunge apparatus housed in a 4°C cold room with 60-70% relative humidity.

### Cryo-EM data acquisition for SMA-cASIC1a

For the resting channel structure at high pH, data were collected on a Titan Krios cryo-electron microscope (ThermoFisher) operated at 300 keV. Images were recorded on a Gatan K3 camera positioned after an energy filter (20 eV slit width) operating in super-resolution mode with a binned pixel size of 0.648 Å. Data were collected with SerialEM (54) and dose-fractionated to 50 frames for a total exposure time of 2-3 s and a total dose of 40-50 e^-^ Å^-2^.

For the desensitized state structure at pH 7.0, data were recorded on a Titan Krios cryo-electron microscope operated at 300 kV and equipped with a spherical aberration corrector. Images were recorded on a Gatan K2 Summit camera in super-resolution mode with a binned pixel size of 1.096 Å. Data were acquired using Leginon (55) and dose-fractionated to 48 frames at 0.15 s per frame for a total exposure time of 7.25 s and a total dose of 50 e^-^ Å^-2^.

### Cryo-EM data processing for SMA-cASIC1a

Images were motion corrected using UCSF MotionCor2 (56) and CTF estimation was performed using Gctf (57). Particles picked using DoG Picker (58) were subjected to reference-free 2D classification in cryoSPARC V2 (59). Following initial classification, an *ab-initio* model was generated in cryoSPARC V2 and used for iterative rounds of 3D classification and refinement in cryoSPARC V2. For the pH 7.0 dataset, per-particle CTF estimation was performed using Gctf. Final reconstructions for both datasets were obtained via non-uniform refinement (C3 symmetry) in cryoSPARC V2.

## ACKNOWLEDGEMENTS

We thank L. Vaskalis for help with figures, H. Owen for manuscript preparation and all Gouaux lab members for their support. Additionally, we thank Polyscope for providing the XIRAN SL 30010 polymer as a gift. This research was supported by the National Institute of Diabetes and Digestive Kidney Diseases (5T32DK007680), and the National Institute of Neurological Disorders and Stroke (5F31NS096782 to N.Y. and 5R01NS038631 to E.G.). Initial electron microscopy work was performed at the Multiscale Microscopy Core at Oregon Health and Science University (OHSU). A portion of this research was performed at the National Center for Cryo-EM Access and Training and the Simons Electron Microscopy Center located at the New York Structural Biology Center, supported by the NIH Common Fund Transformative High Resolution Cryo-Electron Microscopy program (U24GM129539) and by grants from the Simons Foundation (SF349247) and New York State. Subsequent research was supported by NIH grant U24GM129547 and performed at the Pacific Northwest Cryo-EM Center at OHSU and accessed through EMSL (grid.436923.9), a DOE Office of Science User Facility sponsored by the Office of Biological and Environmental Research. Additional support was provided by ARCS Foundation and Tartar Trust fellowships. E.G. is an Investigator with the Howard Hughes Medical Institute.

## AUTHOR CONTRIBUTIONS

N.Y. and E.G. designed the project. N.Y. performed the biochemistry and structural analysis. N.Y wrote the manuscript and all authors edited the manuscript.

## AUTHOR INFORMATION

The authors declare no competing financial interests. Correspondence and requests for material should be addressed to E.G.

## DATA AVAILABILITY

The data that support these findings are available from the corresponding author upon request. The coordinates and associated cryo-EM map for the desensitized SMA-cASIC1a channel at pH 7.0 have been deposited in the Protein Data Bank and Electron Microscopy Data Bank under the accession codes 6VTK and EMD-21380, respectively. The coordinates and associated cryo-EM map for the resting SMA-cASIC1a channel at pH 8.0 have been deposited in the Protein Data Bank and Electron Microscopy Data Bank under the accession codes 6VTL and EMD-21381, respectively.

## SUPPLEMENTARY DATA FIGURE LEGENDS

**Supplementary Data Table 1. Cryo-EM data collection, processing and validation statistics**.

**Supplementary Data Figure 1. Purification of SMA-cASIC1a. a-b**, SDS-PAGE (**a**) and FSEC (**b**) analysis of SMA-solubilized EGFP-cASIC1a particles after size-exclusion chromatography. **c**, Negative stain transmission electron microscopy of SMA-solubilized EGFP-cASIC1a particles.

**Supplementary Data Figure 2. Cryo-EM of SMA-cASIC1a at pH 7.0. a-b**, Representative micrograph (**a**) and 2D classes (**b**) of SMA-cASIC1a at pH 7.0. **c-e**, Local resolution estimation (**c**) angular distribution (**d**) and gold standard FSC resolution estimation (**e**) from final non-uniform refinement in cryoSPARC V2.

**Supplementary Data Figure 3. Cryo-EM data processing for SMA-cASIC1a at pH 7.0.** Data processing strategy for SMA-cASIC1a pH 7.0 dataset.

**Supplementary Data Figure 4. Cryo-EM of SMA-cASIC1a at pH 8.0. a-b**, Representative micrograph (**a**) and 2D classes (**b**) of SMA-cASIC1a at pH 80. **c-e**, Local resolution estimation (**c**) angular distribution (**d**) and gold standard FSC resolution estimation (**e**) from final non-uniform refinement in cryoSPARC V2.

**Supplementary Data Figure 5. Cryo-EM data processing for SMA-cASIC1a at pH 8.0**. Data processing strategy for SMA-cASIC1a pH 8.0 dataset.

**Supplementary Data Figure 6. Polar contacts at the reentrant loop. a**, Polar interactions between the reentrant loop and neighboring residues. **b**, Surface representation of lower TMD showing interfaces between the reentrant loop residues and TM1 and TM2b helices.

