## Supplementary figures and images for "Conserved His-Gly motif of acid-sensing ion channels resides in a reentrant ‘loop’ implicated in gating and ion selectivity"

### Supplemental data figure 1

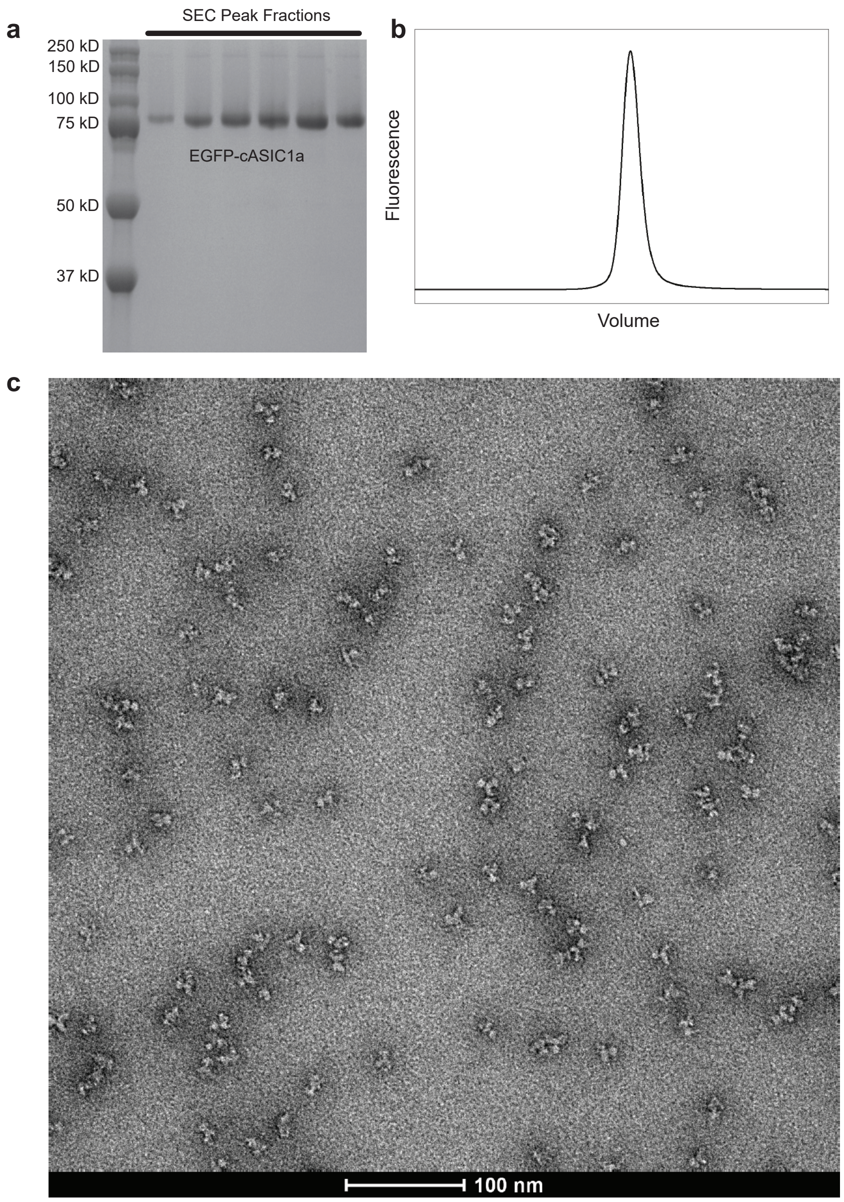


Supplementary data figure 1

### Supplemental data figure 2

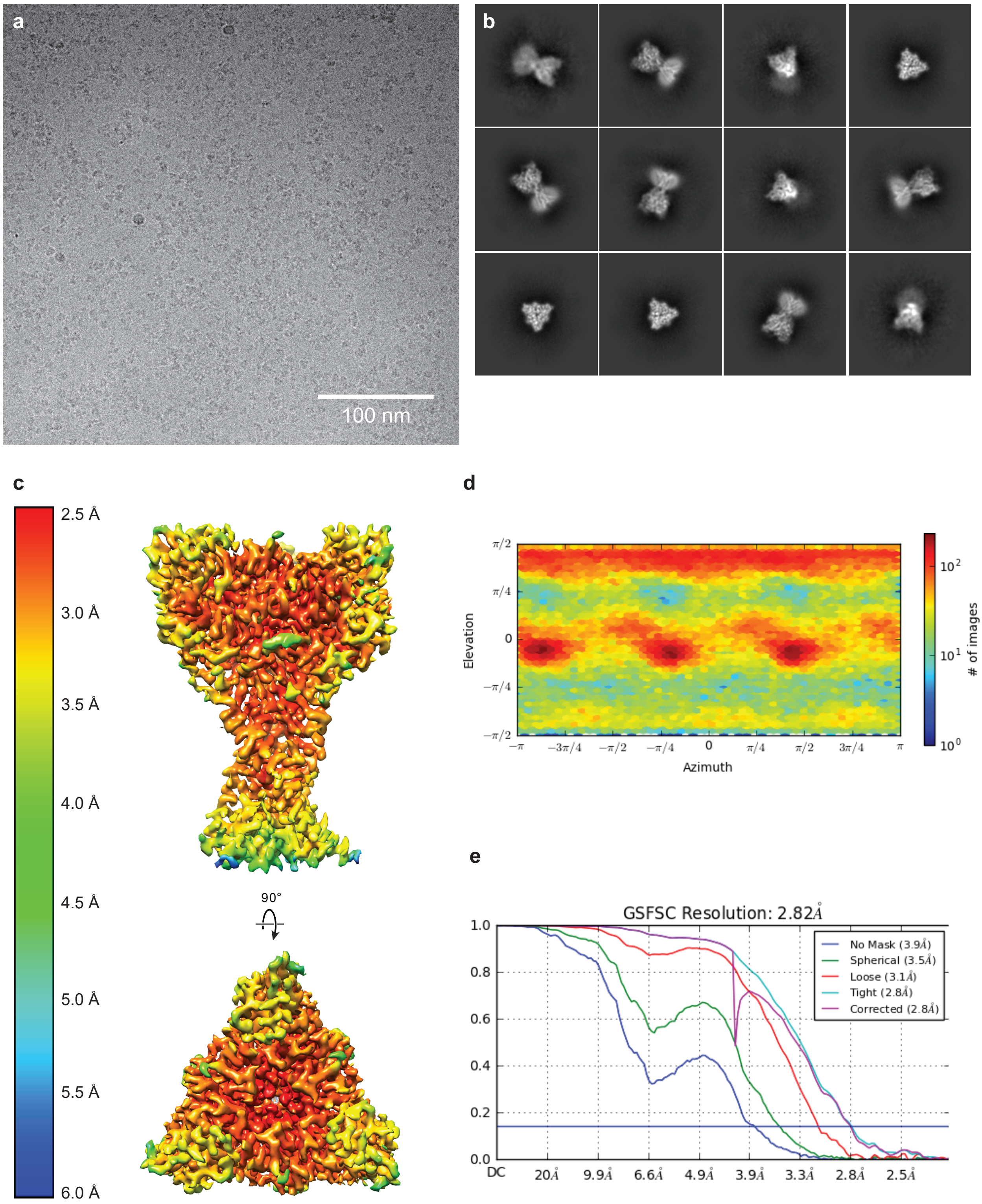


Supplementary data figure 2

### Supplemental data figure 3

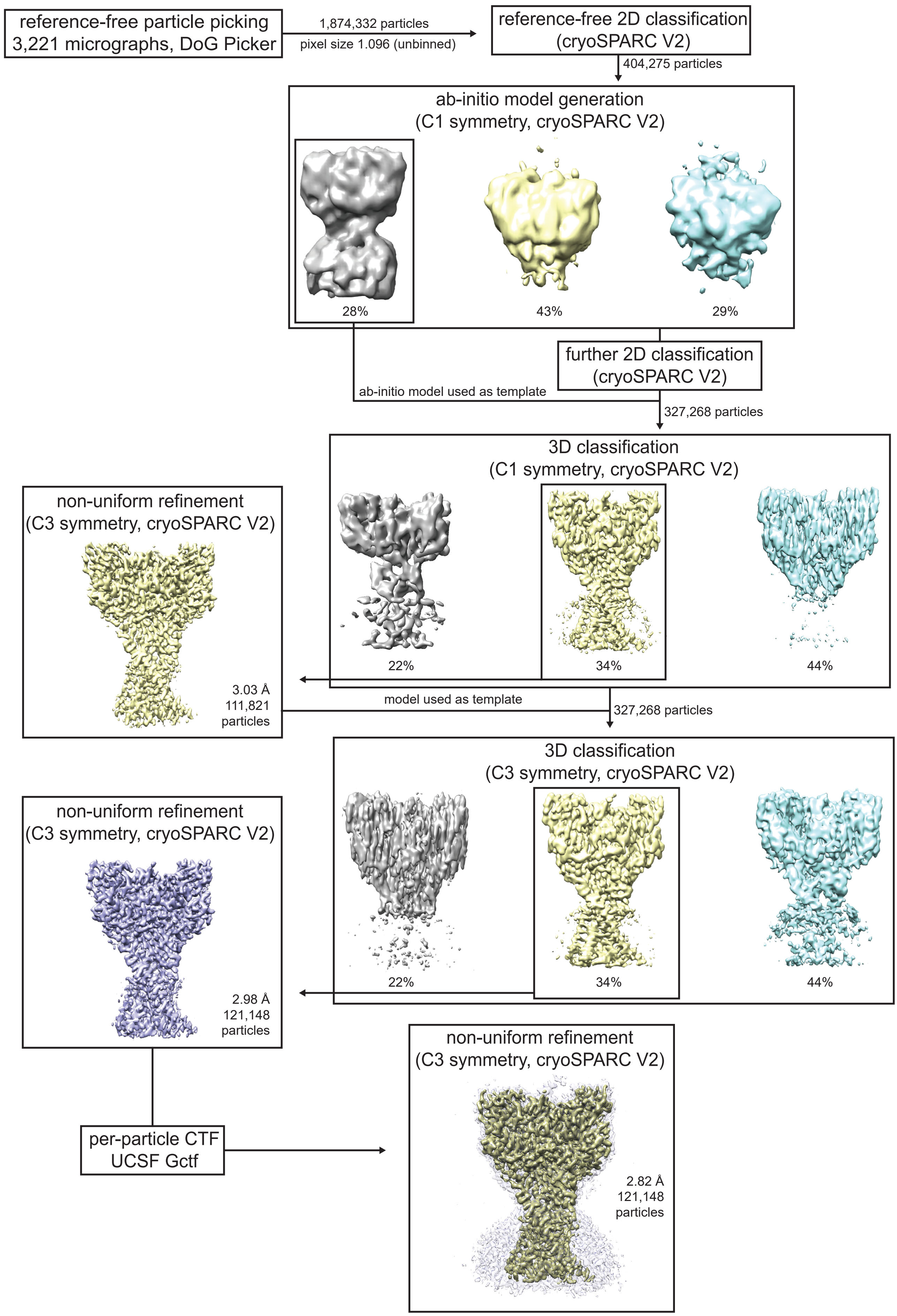


Supplementary data figure 3

### Supplemental data figure 4

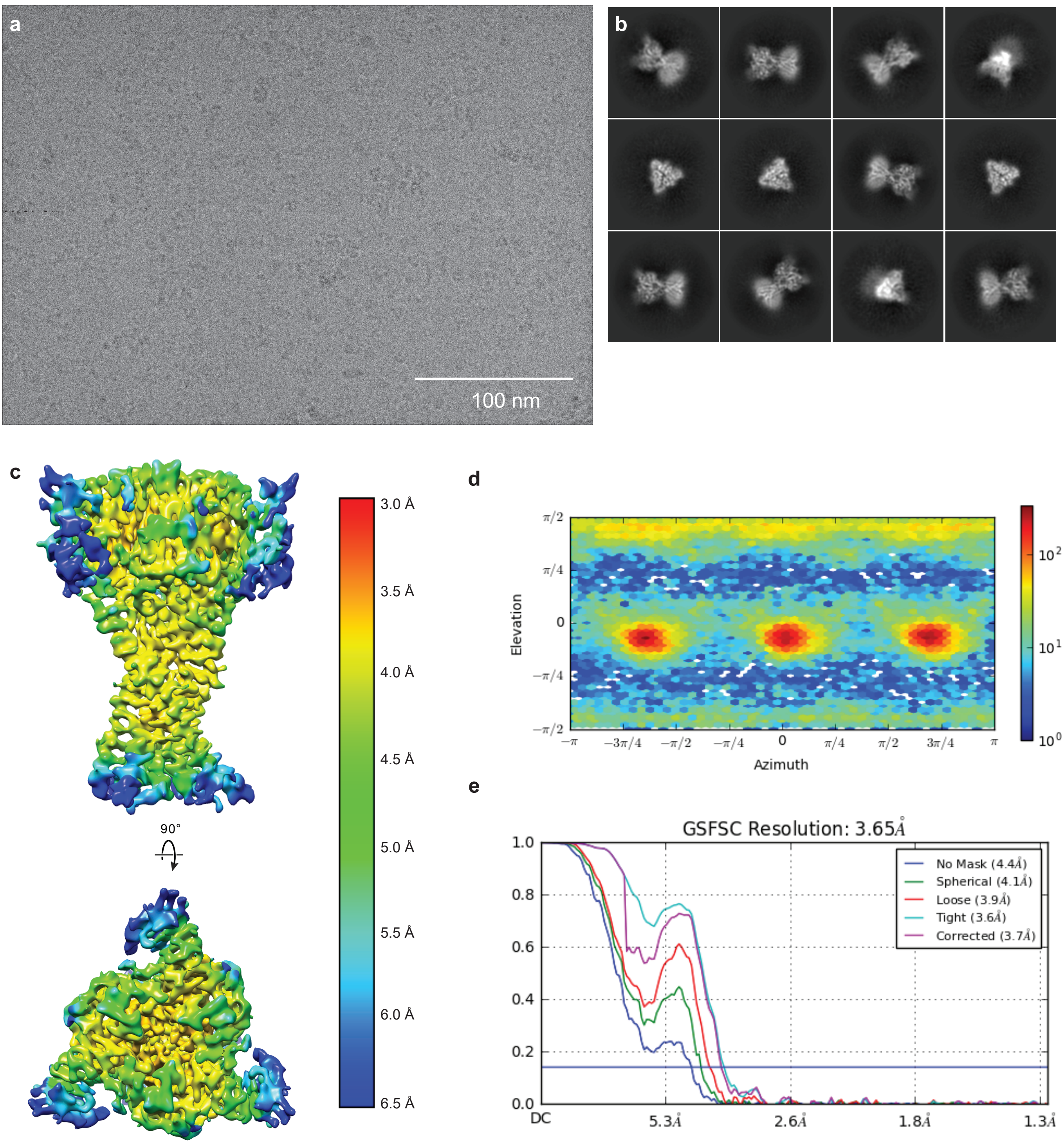


Supplementary data figure 4

### Supplemental data figure 5

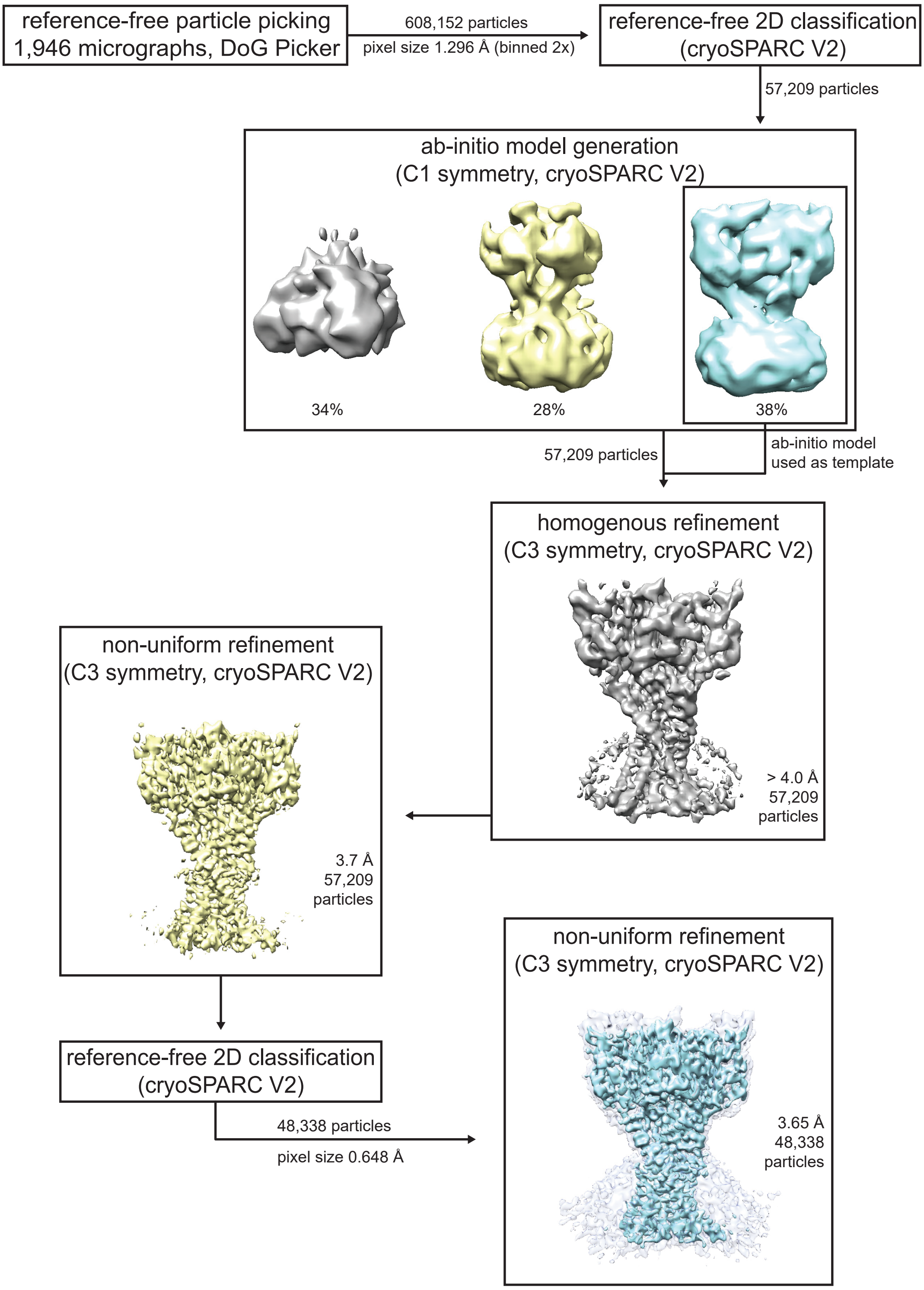


Supplementary data figure 5

### Supplemental data figure 6

**a**

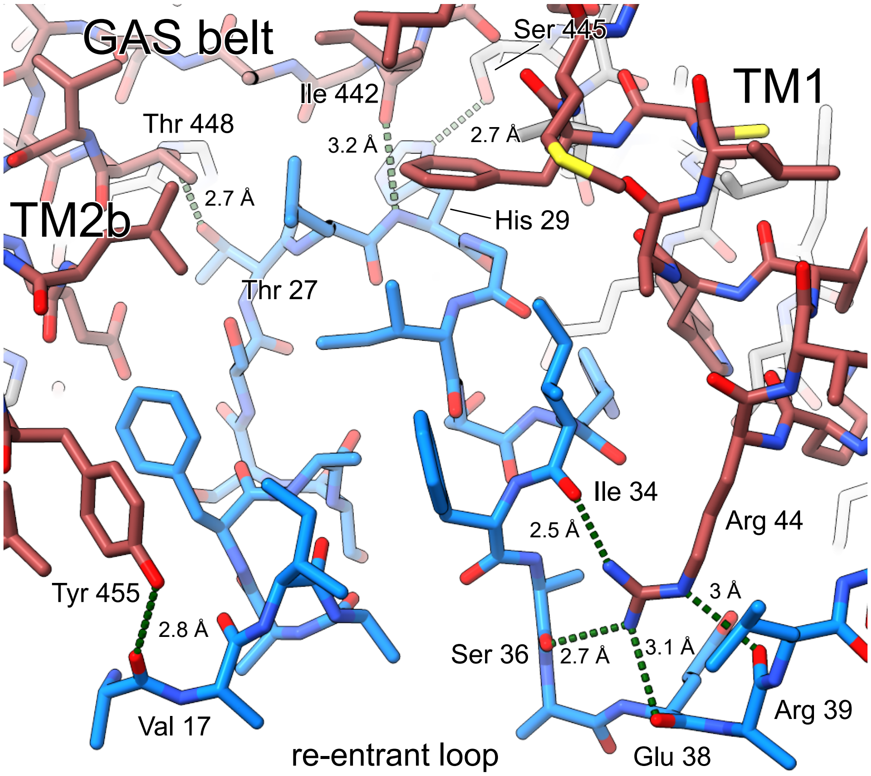


**b**

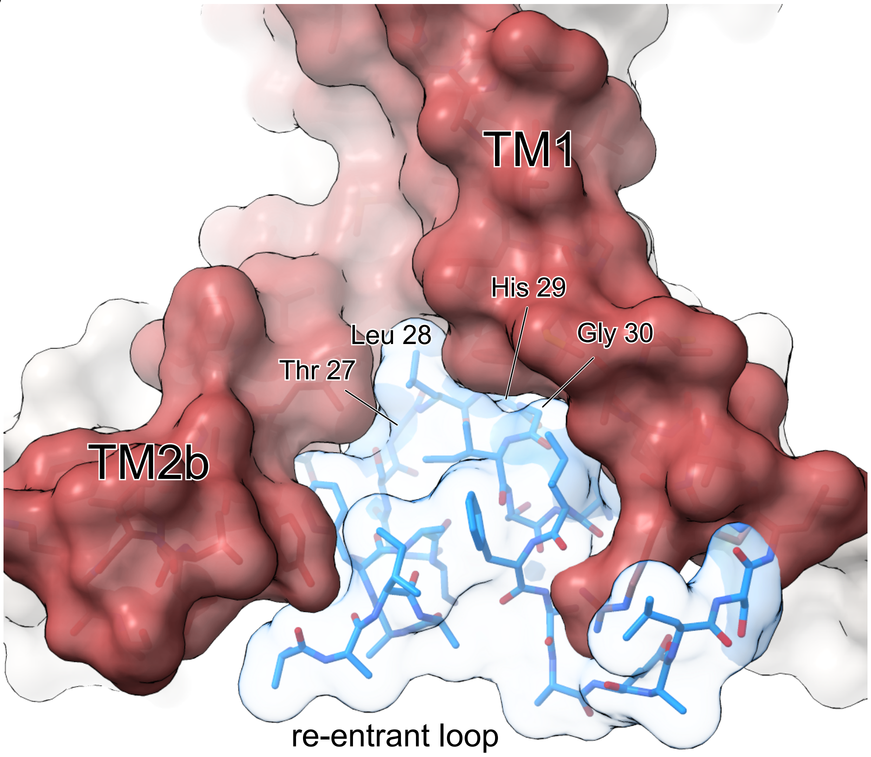


Supplementary data figure 6
