## Supplemental data table 1 for "Conserved His-Gly motif of acid-sensing ion channels resides in a reentrant ‘loop’ implicated in gating and ion selectivity"

**Table 1: Cryo-EM data collection, processing and validation statistics**

| | pH 7.0 SMA-cASIC1a (EMD-21380) (PDB-6VTK) | pH 8.0 SMA-cASIC1a (EMD-21381) (PDB-6VTL) |
|---|---|---|
| ***Data collection/processing*** | | |
| Microscope | Krios (NCCAT) | Krios (PNCC) |
| Camera | K2 | K3 |
| Magnification | 105,000 | 105,000 |
| Voltage (kV) | 300 | 300 |
| Exposure time (s) | 7.25 | 2 |
| Frames (no.) | 48 | 50 |
| Electron exposure (e–/Å$^2$) | 45-50 | 40-50 |
| Defocus range (µm) | -0.8 – -2.6 | -0.8 – -2.6 |
| Pixel size (Å) | 1.096 | 0.648 |
| Initial micrographs (no.) | 3,396 | 1,946 |
| Micrographs used (no.) | 3,221 | 1,946 |
| Particles picked (no.) | 1,874,332 | 608,152 |
| 2D-cleaned particles (no.) | 327,268 | 57,209 |
| ***Final Refinement*** | | |
| Symmetry imposed | C3 | C3 |
| Final particles (no.) | 121,140 | 48,338 |
| Map resolution (Å) | 2.8 | 3.7 |
| FSC threshold | 0.143 | 0.143 |
| ***Model statistics*** | | |
| Initial model used (PBD code) | 4NYK | 5WKU |
| Model resolution (Å) | 3.1 | 4.0 |
| FSC threshold | 0.5 | 0.5 |
| Map sharpening *B* factor (Å$^2$) | 99 | 121 |
| Model composition | | |
| Non-hydrogen atoms | 10686 | 10242 |
| Protein residues | 1338 | 1338 |
| Ligands | 6 | 6 |
| *B* factors (Å) | | |
| Protein | 69.42 | 32.15 |
| Ligand | 89.03 | 124 |
| R.m.s. deviations | | |
| Bond lengths (Å) | 0.005 | 0.004 |
| Bond angles (º) | 0.834 | 0.807 |
| Validation | | |
| MolProbity score | 1.29 | 1.54 |
| Clashscore | 3.10 | 5. 90 |
| Poor rotamers (%) | 0.0 | 0.0 |
| Ramachandran plot | | |
| Favored (%) | 96.85 | 96.62 |
| Allowed (%) | 3.15 | 3.38 |
| Disallowed (%) | 0.0 | 0.0 |
